# Quantifying ERK-activity in response to inhibition of the BRAFV600E-MEK-ERK cascade using mathematical modelling

**DOI:** 10.1101/2021.04.20.440559

**Authors:** Sara Hamis, Yury Kapelyukh, Aileen McLaren, Colin J. Henderson, C. Roland Wolf, Mark A.J. Chaplain

## Abstract

**Background:** Simultaneous inhibition of multiple components of the BRAF-MEK-ERK cascade (vertical inhibition) has become a standard of care for treating BRAF-mutant melanoma. However, the molecular mechanism of how vertical inhibition synergistically suppresses intracellular ERK-activity, and consequently cell proliferation, are yet to be fully elucidated.

**Methods:** We develop a mechanistic mathematical model that describes how the mutant BRAF-inhibitor, dabrafenib, and the MEK-inhibitor, trametinib, affect BRAFV600E-MEK-ERK signalling. The model is based upon a system of chemical reactions that describes cascade signalling dynamics. Using mass action kinetics, the chemical reactions are re-expressed as ordinary differential equations which are parameterised by *in vitro* data and solved numerically to obtain the temporal evolution of cascade component concentrations.

**Results:** The model provides a quantitative method to compute how dabrafenib and trametinib can be used in combination to synergistically inhibit ERK-activity in BRAFV600E-mutant melanoma cells. The model elucidates molecular mechanisms of vertical inhibition of the BRAFV600E-MEK-ERK cascade and delineates how elevated BRAF concentrations generate drug resistance to dabrafenib and trametinib. The computational simulations further suggest that elevated ATP levels could be a factor in drug resistance to dabrafenib.

**Conclusions:** The model can be used to systematically motivate which dabrafenib-trametinib dose-combinations, for treating BRAFV600E-mutated melanoma, warrant experimental investigation.

## Background

Mitogen-activated protein kinase (MAPK) pathways are present in all eukaryotic cells and play a pivotal role in cellular processes linked to proliferation, differentiation and apoptosis^1^. A general MAPK pathway consists of a three-tiered cascade, with a MAPK kinase kinase (MAPKKK) on the top tier, a MAPK kinase (MAPKK) on the middle tier, and a MAPK on the bottom tier. Via sequential kinase activation, signals can propagate through the MAPK cascade in which the activated MAPKKK phosphorylates, and thereby activates the MAPKK, which in turn phosphorylates, and thereby activates, the MAPK.

Several MAPKKK-MAPKK-MAPK cascades are present in eukaryotic cells, but in this study, we focus our attention on one such cascade: the BRAF-MEK-ERK cascade. Mutations in this signalling pathway have been identified as drivers for tumour development and growth, and therefore this cascade is of particular interest in the treatment of cancer. Upon activation, ERK can phosphorylate numerous substrates in the cytoplasm and the nucleus and thereby regulate gene expression directly, by phosphorylating transcription factors such as Elk, Ets and Myc, and indirectly, by acting on substrates that in turn can modify transcription factors^2^. These phosphorylation events, which are controlled by both the amplitude and the duration of ERK activation, are crucial in regulating intracellular processes, including cell proliferation.

In the early 2000s, it was established that BRAF oncogene mutations occur in a majority of melanomas^3^. In melanomas harboring the BRAFV600E mutation, the most common BRAF mutation^4^, the BRAF kinase is constitutively activated and thus signalling through the BRAF-MEK-ERK cascade is hyperactivated and always turned on, which can result in uncontrolled cell proliferation^5,6^. As a consequence, small molecule inhibitors that target the BRAF-MEK-ERK cascade have been developed, with these inhibitors acting to inhibit the enzymatic activity of the kinases in the cascade^7^. Such inhibitors ultimately suppress the amount of activated ERK within a cell and, by extension, the cell’s proliferative abilities. Such drugs have been shown to have profound anti-tumour effects.

*Vertical inhibition* describes a treatment approach in which two or more tiers in a cascade, such as the BRAF-MEK-ERK cascade, are concurrently targeted^8^. In this paper, we specifically investigate drug responses to mono- and combination therapies that include the BRAF-inhibitor dabrafenib and the MEK-inhibitor trametinib. Dabrafenib is an adenosine triphosphate (ATP)-competitive inhibitor with specificity for the MAPKKK BRAFV600E, and trametinib selectively inhibits the MAPKKs MEK1 and MEK2^9^. Trametinib is an allosteric inhibitor as it inhibits the enzymatic activity of MEK1 and MEK2 by binding to a site that is distinct from the ATP-binding site^10^. In 2011, the first selective inhibitor of mutated BRAF, vemurafenib, was approved by the US Food and Drug Administration for treating BRAF-mutant metastatic or unresectable melanoma^11^. Thereafter, selective BRAF-inhibitors replaced the previously used chemotherapies in treating BRAFV600E and BRAFV600K mutant melanoma in clinics, which significantly improved patient survival^12^. However, drug resistance to BRAF-inhibiting monotherapies commonly develops within 6-8 months of treatment^13–15^.

In current treatment protocols, combined BRAF and MEK inhibition is a standard of care for treating BRAFV600E mutant melanoma^13,14,16,17^. Compared to BRAF inhibiting monotherapies, such combination therapies have been shown to delay the onset of drug resistance; in a clinical phase 3 trial, the median progression-free survival was respectively reported to be 5.8 and 9.4 months for dabrafenib mono-treatments and dabrafenib-trametinib combination treatments^14^. This, yet rapid, onset of drug resistance suggests that current treatment regimens create a pressure that selects for drug resistant tumour subclones^15^.

Multiple and diverse mechanisms have been empirically identified as drivers for resistance to BRAF-inhibitors, and one important such mechanism is BRAFV600E amplification^18^. In a multicenter meta-analysis study of clinically observed resistance mechanisms in BRAFV600E-mutated melanoma, quantitative genomic DNA PCR testing showed that BRAFV600E/K amplification was present in 120 out of 132 patient biopsies^19^. Xue et al. moreover experimentally showed that increased BRAFV600E concentrations led to a growth advantage in A375 melanoma cells *in vitro*, when the cells where treated with RAF, MEK or ERK inhibitors^4^. Interestingly, in the absence of these drugs, BRAFV600E amplification did not yield a growth advantage and thus the authors concluded that the magnitude of BRAFV600E amplification required for sustained BRAFV600E-MEK-ERK signalling is drug dependent. Xue et al. referred to this magnitude as the *fitness threshold*. As part of their study, the authors further showed, using patient derived xenografts, that an increased fitness threshold can be obtained via intermittent administration of vertical MAPK inhibition.

Xue et al.’s fitness threshold model can be used to link drug responses with the evolutionary selection of subclones within a tumour, but does not explain why, on an intracellular level, an increase in BRAFV600E concentration leads to a growth advantage *only* in the presence of drugs. However, a recent study by Sale et al. demonstrated that BRAFV600E amplified cells can become addicted to MEK-inhibitors, and that ERK1 or ERK2 activation post drug withdrawal drives G1 cell-cycle arrest, senescence and cell death^20^. This indicates that, during drug withdrawal, BRAFV600E amplified cells have a proliferative disadvantage compared to non-BRAFV600E amplified cells.

In order to further shed light on the effects of vertical inhibition of the BRAFV600E-MEK-ERK cascade at a molecular level, we here develop a mathematical model that mechanistically captures cascade dynamics in the presence of dabrafenib and trametinib. Huang and Ferrell developed the first mathematical model of a *general* MAPK pathway^21^, consisting of a three-tiered cascade in which double phosphorylation of the MAPKKK, MAPKK and MAPK substrates is required for activation. Using mass action kinetics, Huang and Ferrell formulated the MAPK cascade in terms of a system of ordinary differential equations, which they solved numerically to simulate signalling dynamics within the cascade. Notably, the authors showed that the structure of the MAPK cascade yields ultra-sensitive signalling dynamics. In this study, we modify Huang and Ferrell’s model to capture cellular signalling dynamics in a *specific* MAPK pathway: the BRAF-MEK-ERK cascade in BRAFV600E mutant melanoma. As an extension to Huang and Ferrell’s model, we introduce ATP-dependent substrate phosphorylation, and molecular drug actions of dabrafenib and trametinib.

## Methods

We develop a mathematical model that simulates the temporal BRAF-MEK-ERK signalling dynamics in BRAFV600E mutant melanoma. In order to do this, we first map out the structure of the BRAF-MEK-ERK cascade, and identify the drug actions of dabrafenib and trametinib, that respectively inhibit enzymatic BRAF and MEK activity. From the cascade structure and drug actions, we formulate a system of chemical reactions which captures intracellular signalling dynamics. Using mass action kinetics, the system of chemical reactions is thereafter re-formulated as a system of ordinary differential equations (ODEs) which is solved numerically in order to simulate the temporal evolution of signalling molecule concentrations. A model output of particular importance is the concentration of activated ERK over time, as the ultimate goal of vertical inhibition of the BRAFV600E-MEK-ERK pathway is to suppress cells’ activated ERK levels and, by extension, their proliferative abilities. All model parameters (*i*.*e*., kinetic constants and total intracellular protein concentrations) are obtained from data available in the literature^21–29^.

### The BRAF-MEK-ERK cascade

We modify Huang and Ferrel’s cascade model^21^ to specifically capture the signalling dynamics of the BRAF-MEK-ERK cascade in BRAFV600E mutant melanoma. Both Huang and Ferrell’s general MAPKKK-MAPKK-MAPK cascades and our problem-specific cascade are respectively illustrated in Figures 1a and b. Here, and throughout this work, singly and doubly phosphorylated proteins are respectively denoted by the prefixes p and pp. Note that double phosphorylation is required for substrate activation, and that phosphatases (phosphs) dephosphorylate singly- and doubly phosphorylated MAPKK/MEK and MAPK/ERK (Figure 1). In our model, BRAF phosphorylates MEK and once MEK has been doubly phosphorylated, it in turn phosphorylates ERK, which becomes activated when doubly phosphorylated. As BRAF is constitutively activated in BRAFV600E-mutant melanoma^6^, we have reduced the top tier in our modified cascade to contain only activated BRAF (Figure 1b). Since the model developed in this study will include the drug actions of dabrafenib, which is an ATP-competitive inhibitor, and trametinib, which is an allosteric ATP inhibitor, we extend Huang and Ferrell’s model to include ATP-dependent substrate phosphorylation. Thus, in our model, when a substrate and an ATP molecule bind to an enzyme (in the absence of drugs), the ATP molecule donates a phosphate group to the substrate, and the ATP converts to adenosine diphosphate (ADP) after the metabolic process (Figure 1b).

**Figure 1:**
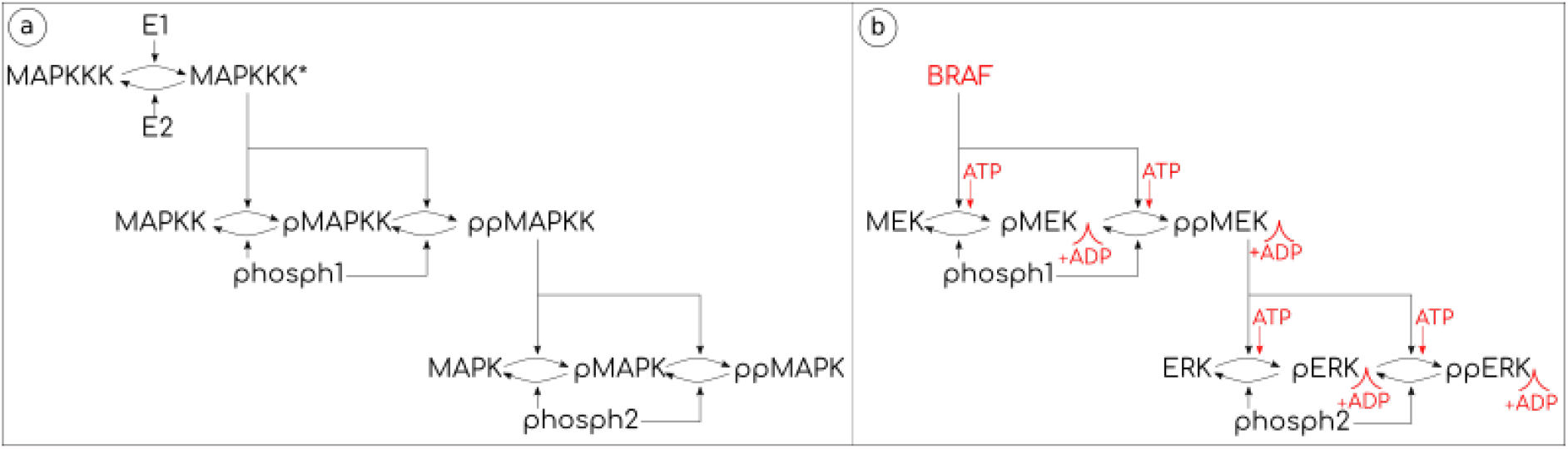
Cascade models. (a) The three-tiered general MAPKKK-MAPKK-MAPK cascade of Huang and Ferrell^21^. The star (*) denotes activated MAPKKK, and E1 and E2 are unspecified enzymes that respectively activate and inactivate MAPKKK. The prefixes p and pp denote singly and doubly phosphorylated proteins, and phosph1 and phosph2 are phosphatases that dephosphorylate MAPKK and MAPK respectively. (b) Our modified BRAF-MEK-ERK cascade in BRAFV600E mutant melanoma, in which a specific MAPKKK (BRAF), MAPKK (MEK) and MAPK (ERK) are considered. BRAF is constitutively activated, and therefore the top cascade tier is reduced to a single node, activated BRAF. The modified cascade incorporates ATP-dependent substrate phosphorylation at the sites shown. Differences in the cascade structure between Huang and Ferrell’s original model, and our model, are marked in red in (b).

### BRAF and MEK inhibitors

We model the pharmacodynamic actions of the two drugs dabrafenib and trametinib. Dabrafenib (DBF) is a potent and selective BRAF inhibitor, which is ATP-competitive^30^. As illustrated schematically in Figure 2 (top), DBF can thus bind to the ATP site of a BRAF molecule, and thereby inhibit BRAF-ATP reactions and the downstream phosphorylation of MEK and pMEK. Trametinib is classified as an allosteric MEK1/2 inhibitor, as it binds to a binding site that is distinct from, but adjacent to, the ATP binding site on the MEK molecule^31^. This is illustrated schematically in Figure 2 (bottom). MEK-bound trametinib prevents MEK from phosphorylating the substrates ERK and pERK.

**Figure 2:**
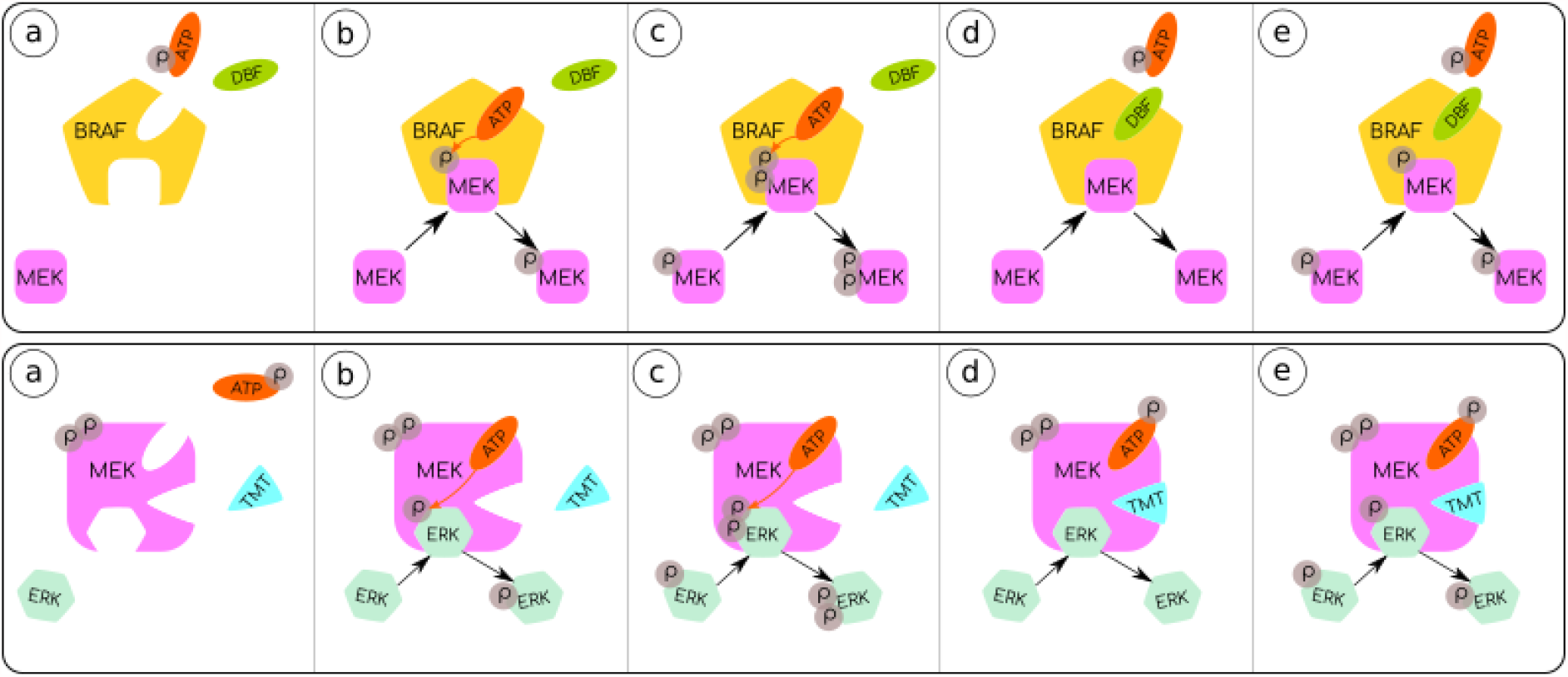
Schematics of the mathematical models that describe dabrafenib (DBF) and trametinib (TMT) drug actions. Top: (a) We consider two BRAF binding sites: one which can bind MEK and one which can bind ATP or dabrafenib. BRAF-bound ATP is necessary for the phosphorylation of MEK. (b) If ATP and MEK are bound to BRAF, then MEK is phosphorylated to the singly phosphorylated form pMEK. (c) If ATP and pMEK are bound to BRAF, then pMEK is phosphorylated to the doubly phosphorylated form ppMEK. (d) and (e) BRAF-bound dabrafenib prevents ATP from binding to BRAF, and thereby inhibits the phosphorylation of the substrates MEK and pMEK. Bottom: (a) We consider three MEK binding sites: one which can bind ERK, one which can bind ATP and one which can bind trametinib. ppMEK-bound ATP is necessary for the phosphorylation of ERK. (b) If ERK and ATP, but not TMT, are bound to ppMEK, then ERK is phosphorylated to the singly phosphorylated form pERK. (c) If pERK and ATP, but not TMT, are bound to ppMEK, then pERK is phosphorylated to the doubly phosphorylated form ppERK. (d) and (e) If TMT is bound to ppMEK, the phosphorylation of the substrates ERK and pERK is inhibited.

### The system of reactions

By combining the cascade illustrated in Figure 1b, with the drug actions that are schematically summarised in Figure 2, we formulate a system of chemical reactions that describes the signalling dynamics in the BRAFV600E-MEK-ERK cascade subjected to drugs. This results in a system of 36 chemical reactions (R.1-R.36), that are outlined in Figure 3, where the order of the binding events is considered.

**Figure 3:**
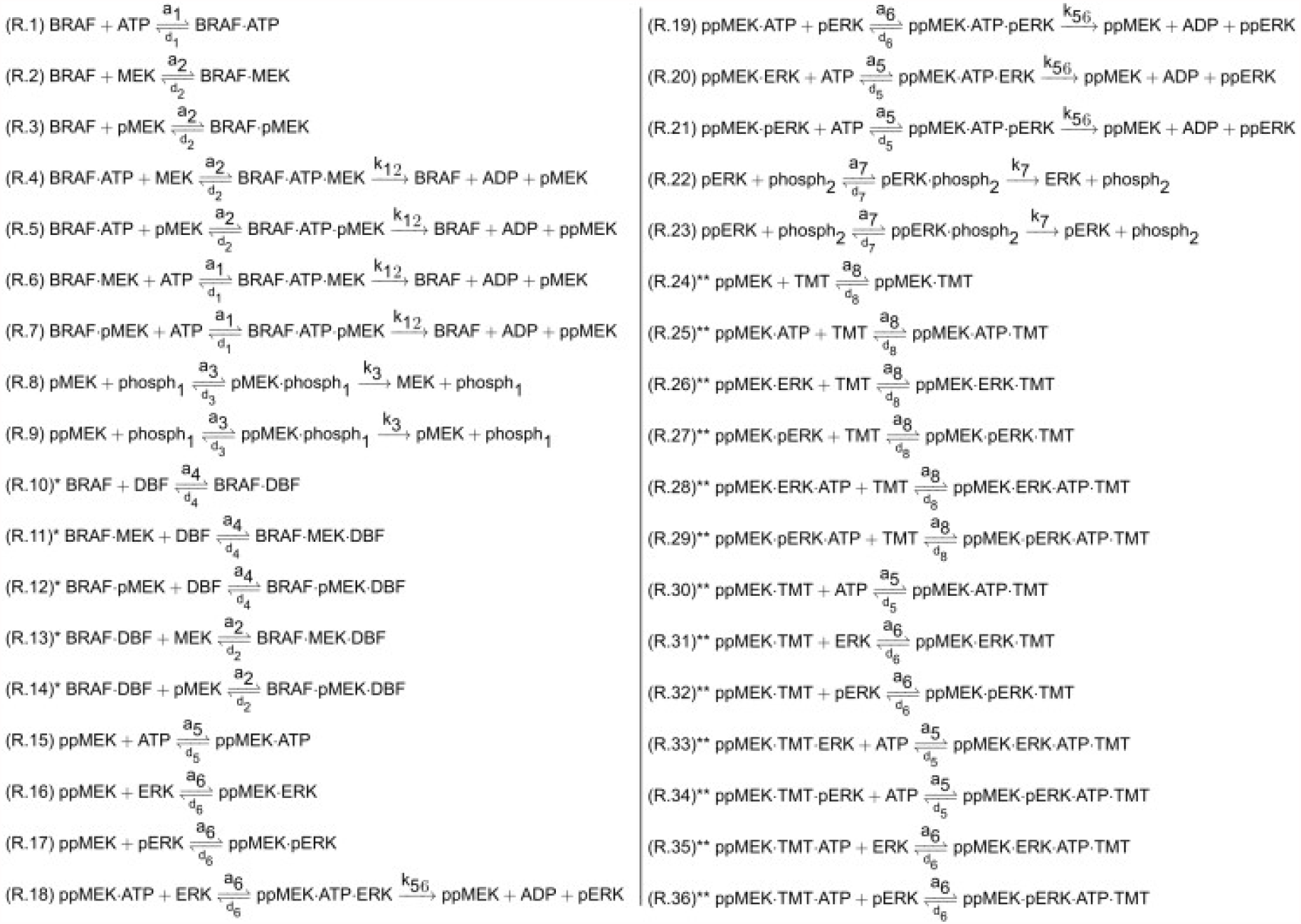
System of reactions (R.1-R.36) describing the signalling dynamics in the BRAFV600E-MEK-ERK cascade subjected to the drugs dabrafenib (DBF) and trametinib (TMT). The dot-notation (·) denotes complexes of two or more molecules. Reactions involving dabrafenib and trametinib are respectively marked by single (*) and double (**) stars and can be omitted when simulating the system in the absence of these inhibitors.

### The system of ordinary differential equations

We use the law of mass action to re-express the system of reactions (R.1-R.36) as a system of ODEs. In the system of ODEs, the dependent variables are the concentrations of the signalling molecules that occur in (R.1-R.36), and the independent variable is time. In short, the law of mass action says that the rate of a chemical reaction is proportional to the product of the masses of the reactants. Thus, the first reaction, R.1, which includes the molecules BRAF, ATP and BRAF•ATP, yields the following contributions to the system of ODEs,

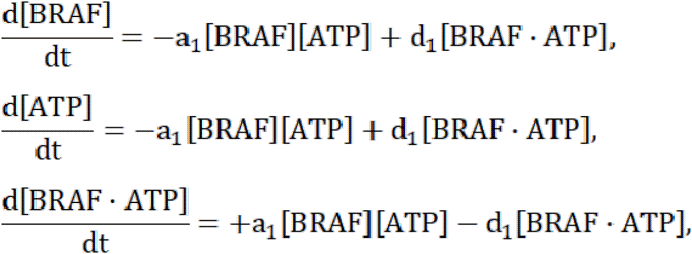

where concentrations are denoted by square brackets surrounding the signalling molecule of interest, and *t* denotes time. Note that the right-hand sides of the above equations are subsequently extended to include more equations derived from other reactions in the system of reactions outlined in Figure 3. The full system of ODEs (O.1-O.36), is available in the Supplementary Material (SM1). In this work, we combine (O.1-O.36) with a set of system conservation laws (C.1-C.7 in SM1) to formulate a system of differential algebraic equations that we solve numerically to obtain the temporal evolution of the system molecule concentrations. The conservation laws ensure that the total concentrations of the kinases BRAF, MEK and ERK (in free, bound, non-phosphorylated, singly phosphorylated and doubly phosphorylated form), phosphatases (in free and bound form) and drugs (in free and bound form) are conserved within the system.

### Model implementation

We implement and solve the system of ODEs (O.1-O.36), with conservation laws (C.1-C.7) *in silico*, in order to simulate the signalling dynamics in the BRAF-MEK-ERK cascade. The *in silico* model, which is developed using the programing language and computing environment MATLAB, is available on the code hosting platform GitHub. Instructions on how to access, run and modify the code are available in the Supplementary Material (SM2).

### Model parameters

The system of reactions (R.1-R.36) includes eight forward rate constants a_1_, a_2_, …,a_8_, eight reverse rate constants d_1_, d_2_, …, d_8_, and four catalytic rate constants k_1_,_2_, k_3_, k_5_,_6_, k_7_. The values of these constants are listed in Table ST1 in the Supplementary Material (SM3). Huang and Ferrell^21^ argued that, when computing steady-state enzyme levels, it is not the individual rate constants, but rather the Michaelis constants, K_mi_=(d_i_+k_i_)/a_i_, that are important. We therefore set all forward rate constants a_i_ to be the same so that a_j_=a_1_ for all j=2,3,..,8. We then use data available in the literature to set the parameter values for a_1_, the eight reverse rate constants and the four catalytic rate constants, as is outlined in the Supplementary Material (SM4). The model initial condition, *i*.*e*., a vector that includes the molecule concentrations at the start of the simulation, is also obtained from data in the literature^21,28^. Initial cascade component concentrations are listed in Table ST2 in the Supplementary Material (SM3).

## Results

Using the *in silico* model that we have developed, the temporal evolution of the concentration of all signalling components appearing in the system of reactions (R.1- R.36) are computed. In this study, our main model output of interest is activated ERK. For ease of presentation, we define a measure activated ERK, as a time-dependent fraction,

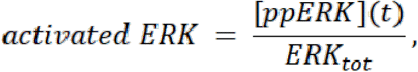

where *ERK*_*tot*_ denotes the total amount of ERK in the system, such that *ERK*_*tot*_*=*[ERK](0), as listed in Table ST2 in the Supplementary Material (SM3).

### Monotherapy results

We initially investigate how activated ERK levels within a cell change in response to dabrafenib or trametinib monotherapy. The temporal evolution of activated ERK in response to these monotherapies are shown in Figures 4a and b respectively.Activated ERK levels at specific time points are plotted over intracellular concentrations of BRAF (left panel) and ATP (right panel) for different doses of dabrafenib (Figure 4c) and trametinib (Figure 4d).

**Figure 4:**
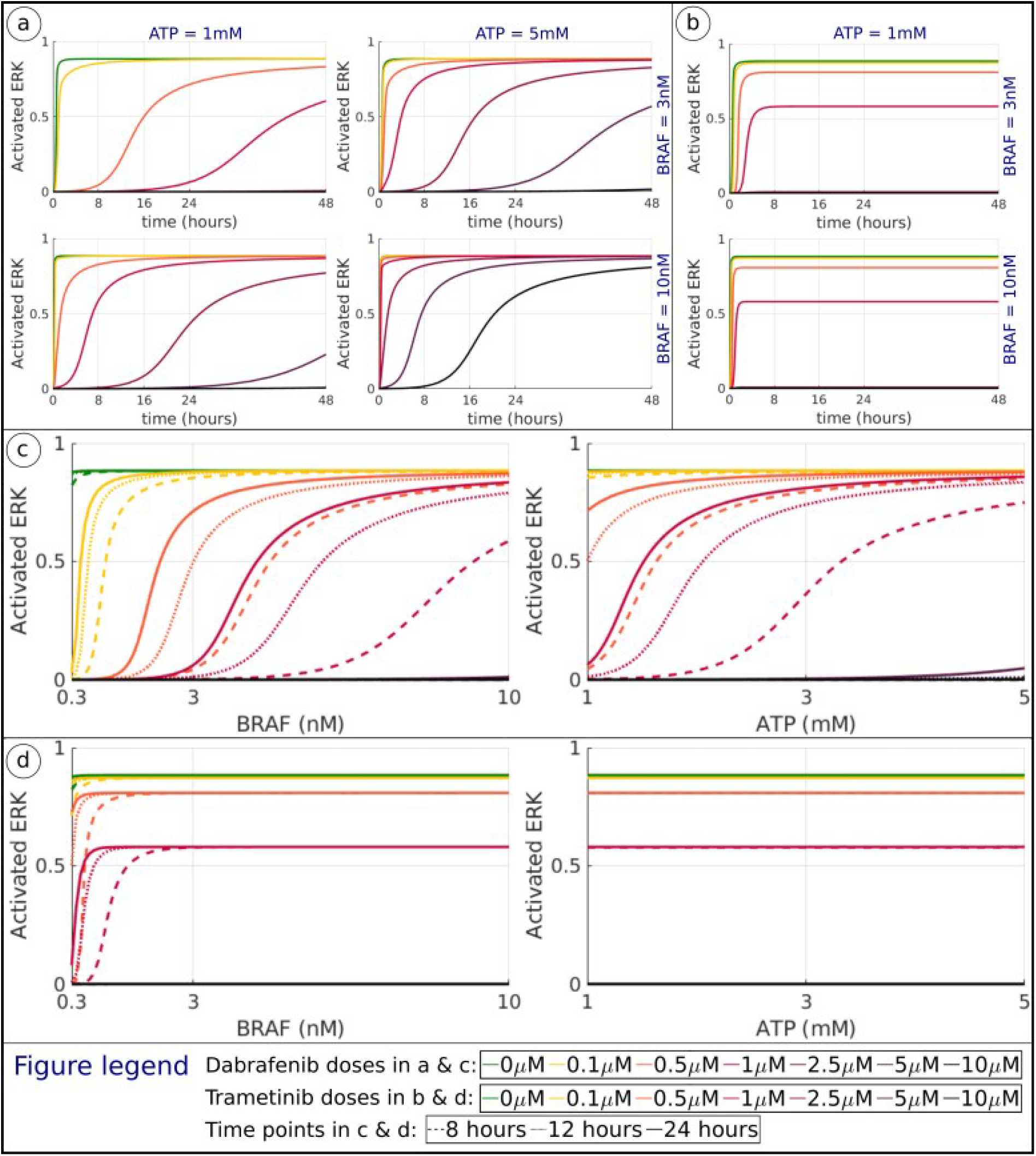
Plots showing activated ERK levels (as defined in Equation 2) in response to monotherapies of dabrafenib or trametinib, where the drug doses are indicated in the legend at the bottom of the figure. (a) The temporal evolution of activated ERK. Results are shown for different intracellular concentrations of BRAF, ATP and dabrafenib (DBF). (b) The temporal evolution of activated ERK. Results are shown for different intracellular concentrations of BRAF and trametinib (TMT). (c,d) Activated ERK levels are plotted over intercellular BRAF concentrations, where ATP concentrations are fixed at the baseline value 1mM (left). Activated ERK levels plotted over intercellular ATP concentrations, where BRAF concentrations are fixed at the baseline value 3nM (right). Results at 8, 16, and 24 hours are shown in dashed, dotted and solid lines respectively for different doses of dabrafenib (c) and trametinib (d).

The simulation data show that the modelled dabrafenib treatment responses are highly dynamic in time, and dependent on both intracellular BRAF and ATP concentrations (Figures 4a and 4c). These data also suggest that increased concentrations of BRAF and ATP make the cells less sensitive to dabrafenib. Further, these data show that the activated ERK levels eventually reach the same steady state value for several dabrafenib-BRAF-ATP concentration combinations (see also Supplementary Material, SM5). In order for a cell to progress from the G1 to the S phase of the cell cycle, sustained ERK-activity throughout the G1 phase is required, where ERK downregulates several anti-proliferative genes until S phase entry^32,33^. This suggests that, if ERK-activity can be suppressed for the duration of the G1 cell cycle phase, further cell cycle progression, and thus cell proliferation, can be inhibited.

In response to different trametinib doses, activated ERK levels reach distinct steady-state levels (Figure 4b). Moreover, trametinib treatment responses at time points after 8 hours are only sensitive to BRAF variations for low BRAF concentrations (under 3nM) and the efficacy of trametinib treatments is not sensitive to variations in ATP within the range of concentrations tested (Figure 4d).

### Combination therapy results

We next evaluate the effect of combination therapies involving dabrafenib and trametinib on treatment outcome (Figure 5). When dabrafenib concentrations are zero (*i*.*e*., for trametinib monotherapies) the activated ERK levels do not vary between 8 to 24 hours. However, when trametinib concentrations are zero (*i*.*e*., for dabrafenib monotherapies) the activated ERK levels do vary between 8 to 24 hours and, as a consequence, so does dabrafenib-trametinib combination therapies (Figure 5a-c).

**Figure 5:**
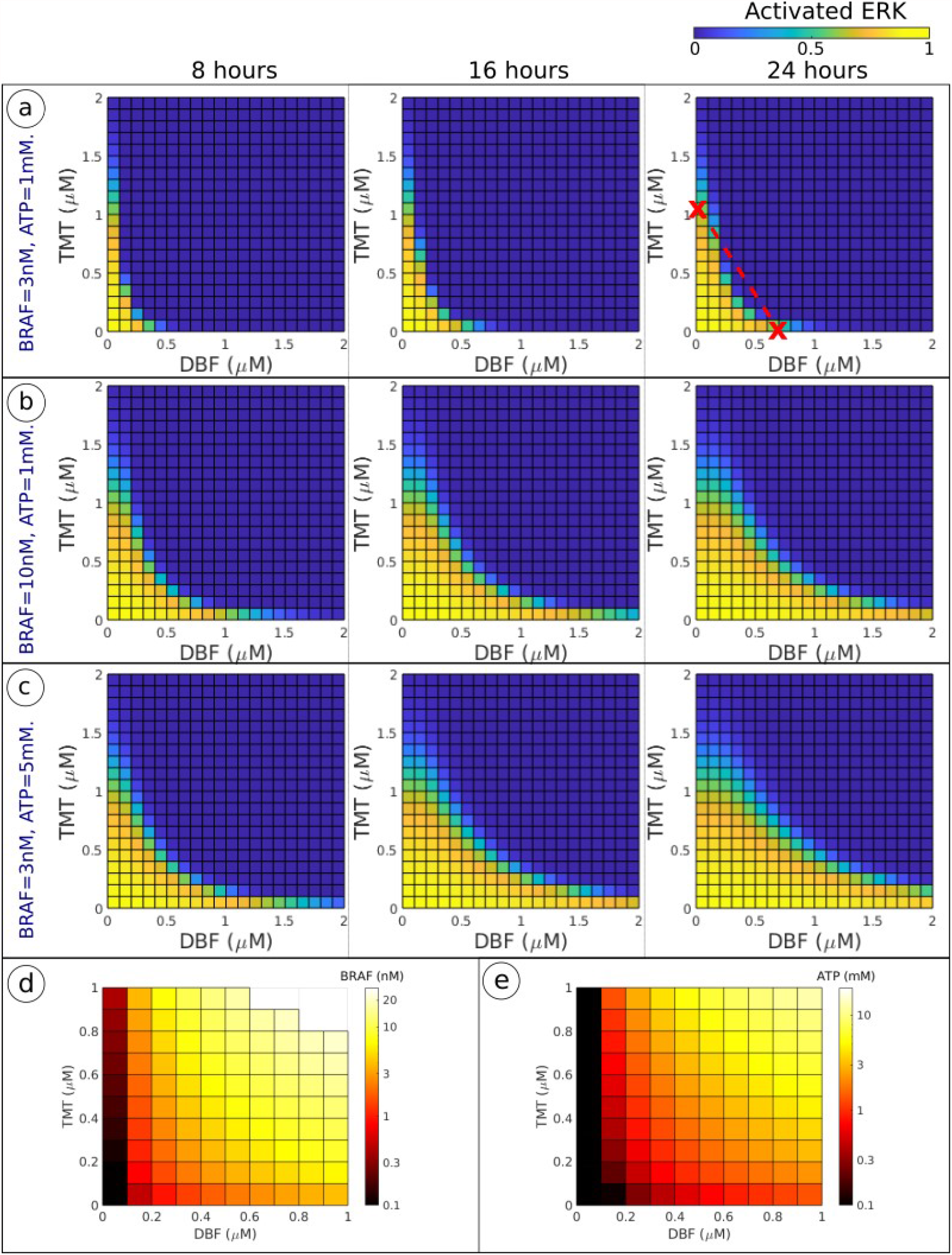
Heatmaps showing treatment responses to dabrafenib-trametinib (DBF-TMT) combination therapies. For each square in the heatmap, the horizontal axis of the intersection denotes the dabrafenib dose, and the vertical axis denotes the trametinib dose. (a,b,c) The fraction of activated ERK (as defined in Equation 2) at 8, 16 and 24 hours are shown in the left, middle and right columns respectively. BRAF and ATP concentrations vary between (a), (b) and (c). The right panel in (a) includes an isobole that connects the two monotherapy drug doses that yield an activated ERK level of 0.5 with a straight red line. Drug combinations below this line that yield activated ERK levels of 0.5 or less are categorised as synergistic. (d,e) The minimum BRAF (d) and ATP (e) concentrations that are required to yield an activated ERK level over 0.5, at 24 hours are shown. In (d), the ATP concentration is fixed at 1 mM. In (e), the BRAF concentration is fixed at 3 nM. For DBF-TMT combination doses where the activated ERK does not surpass 0.5 for BRAF concentrations under 20 nM, the squares in the heatmap in (d) are not coloured.

In order to determine whether there is dabrafenib-trametinib synergism we included an isobole that connects the two monotherapy doses that yield an activated ERK level of 0.5 (Figure 5a). Since drug combinations below this line can achieve activated ERK levels lower than 0.5, the drug combination is categorised as synergistic^34^.

In Figures 5b and c, the BRAF and ATP levels have been increased from the baseline parameter values. By comparing Figure 5a, with Figures 5b and c, we can see that activated ERK levels increase with increased BRAF and ATP levels. Our mathematical model therefore predicts that increased intracellular BRAF and ATP concentrations will generate resistance to dabrafenib-trametinib combination therapies.

The influence of BRAF concentration on dabrafenib-trametinib efficacy is also reflected in Figure 5d. We see from these results the minimum BRAF concentration that is required to achieve an activated ERK level of 0.5 at 24 hours (when ATP is fixed at the baseline value 1 mM). Similarly, the minimum ATP concentration that is required to achieve an activated ERK level of 0.5 at 24 hours (when BRAF is fixed at the baseline value 3 nM) is visualised in Figure 5e.

## Discussion

Although targeted anticancer drugs can generate patient responses in tumours previously intractable to treatment, there remain major issues in their clinical use. In particular, tumour regrowth due to rapid onset of drug resistance. A number of mechanisms of drug resistance have been described for example for drugs targeted at mutant BRAF in the MAPK pathway. Many of these involve the modulation of components of the pathway *e*.*g*., by mutation or gene amplification. This results in the sustained activation of ERK in the presence of drug. As a consequence, drugs targeted at multiple components of the same pathway have been used in combination *e*.*g*., BRAF and MEK inhibitors. Although this has resulted in some patient benefit, this has not resolved the problem and major challenges still remain. There is currently great interest in further developing the complexity of combination treatments. In the case of the BRAF pathway, for example, by the addition of an inhibitor of ERK^4^.

Increasing treatment complexity however raises some major pharmacological challenges, including which drugs to use in combination and what doses to use in order to keep toxicity at an acceptable level. At clinically administered doses, BRAF and MEK inhibitors lead to a number of side-effects that are generally categorised to be non-life threatening and safe when adequately monitored^35,36^. Common adverse events induced by dabrafenib include hyperkeratosis, headache, arthralgia and pyrexia, whilst trametinib commonly induces rash, diarrhoea, fatigue, peripheral oedema and acneiform dermatitis. Dabrafenib-trametinib combination regimens have not been reported to yield any new adverse events. However, fever, chills, fatigue, diarrhoea, hypertension and vomiting are more frequently observed in response to combination therapies than in response to the corresponding monotherapies.

Contrariwise, cutaneous adverse events, such as squamous cell carcinoma and skin papilloma, occur less frequently in response to dabrafenib-trametinib combination therapies than in response to dabrafenib monotherapies^35^.

The vast number of drug combinations and dosing regimens which could be evaluated make it impossible to test all these either in the laboratory or in the clinic. Informative *in silico* pharmacological models are therefore needed.

Several research groups have analysed and modified Huang and Ferrell’s MAPK cascade model since it was presented in 1996^21^, as is summarised in a review by Orton *et al*.^2^. Mathematical analysis and computational simulation of the MAPK cascade model has explained how the dual phosphorylation and dephosphorylation events mechanistically give rise to the ultrasensitivity of ERK activation^37,38^ and bistability^39^. Integrated *in vitro*-mathematical work has further demonstrated switch-like (on or off) ERK-activity in single cells in Xenopus oocytes subjected to progesterone^40^. Moreover, MAPK cascade models of various complexities have been presented and investigated in the literature, where these have included *e*.*g*., the internalisation and downstream signalling activity of the epidermal growth factor receptor^41,42^.

To our knowledge, the mathematical model presented in this paper is the first MAPK cascade model to *(i)* specifically capture signalling dynamics in the BRAFV600E-MEK-ERK cascade, *(ii)* explicitly include ATP-dependent substrate phosphorylation, and *(iii)* include mechanistic drug actions of dabrafenib and trametinib, where the model parameters involved in *(i-iii)* are obtained from published *in vitro* data.

In this paper, we have developed a mathematical model that captures signalling dynamics in the BRAFV600E-MEK-ERK cascade, in response to dabrafenib and trametinib treatments. Our *in silico* model can be used to quantify the temporal evolution of system molecule concentrations and, as a consequence, ERK-activity. The model predicts that increased cellular BRAF and ATP concentrations will result in reduced sensitivity to dabrafenib-trametinib combination therapies. The prediction that ATP levels may influence drug sensitivity introduces an important variable to take into consideration when evaluating treatment responses. Intratumoural heterogeneity generated by both spatio-temporal factors (such as varying ATP levels within a tumour) and phenotypic factors (such as differences in BRAF amplification between cells) has been recognised to fuel drug resistance and complicate the design of treatment strategies^43,44^. Our finding that increased BRAF concentration is associated with drug resistance is furthermore consistent with experimental data^4^.

Whilst simulation of the ERK activation level at different concentrations of dabrafenib and 3 nM BRAF closely followed that observed in patients^45^, an increase of the BRAF level to 10 nM in our model, similar to that found in drug resistant patient derived xenografts^4^, predicted marked resistance of the ERK activation to treatment with dabrafenib. This is shown in Figure 6, where simulation results are superimposed over clinical data obtained from a study in which phosphorylated ERK levels were measured in BRAFV600-positive melanoma biopsy samples before, and five days after, dabrafenib monotherapy administration where doses between 70 and 200 mg were administered twice daily^45^. The recommended dabrafenib dose for unresectable or metastatic BRAFV600E positive melanoma is 150 mg twice daily^6^.

**Figure 6:**
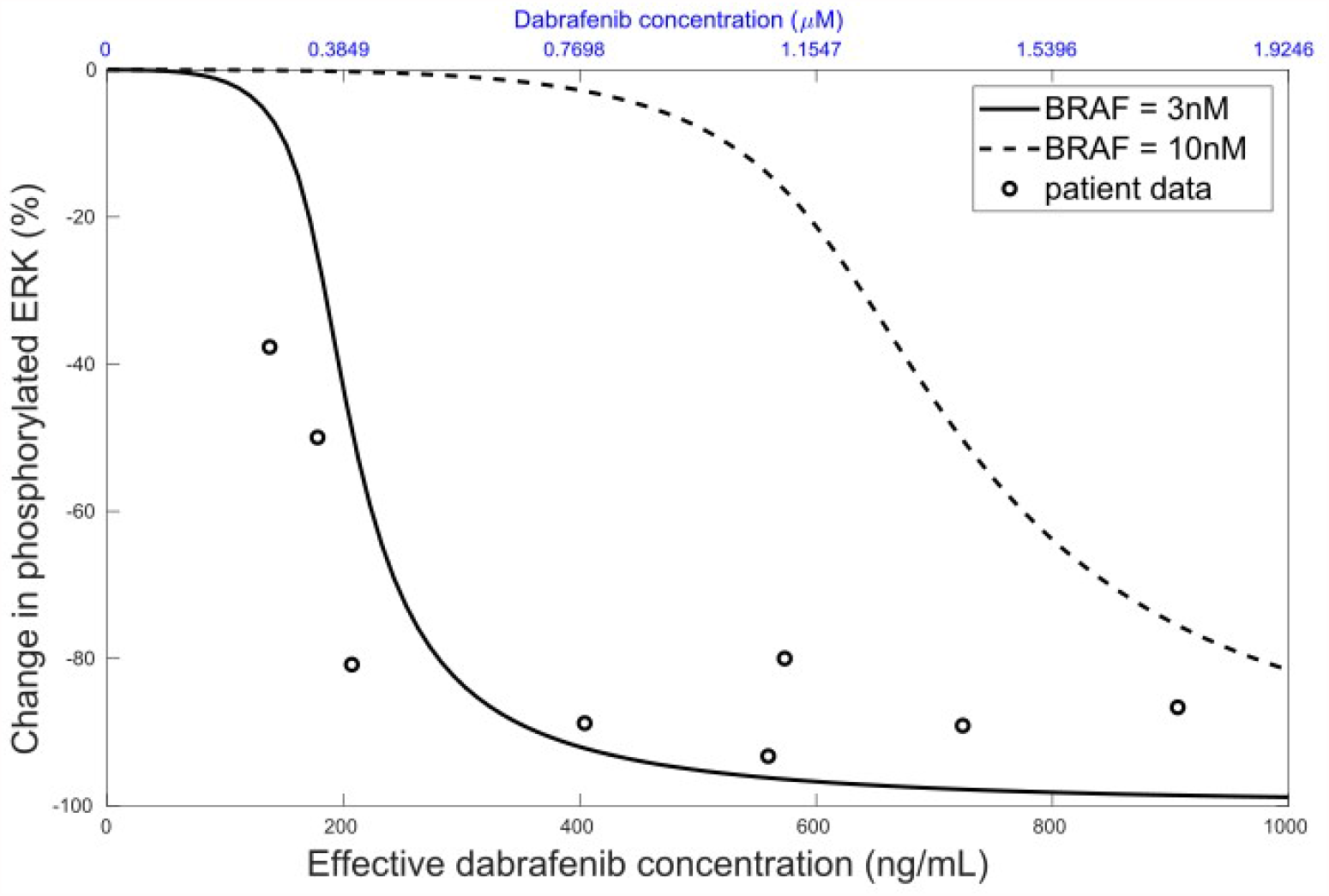
Plots showing changes in phosphorylated ERK ([pERK]+[ppERK]) from the baseline (no drug) level. Simulation results at 8 hours, when the ATP concentration is fixed at 1mM, are shown for BRAF concentrations of 3 and 10nM. The circles show patient data from a clinical study by Falchook *et al*.^45^ where dabrafenib was administered twice daily at doses between 70 and 200 mg. Dabrafenib concentrations are shown µM (used in other simulations results in this article) along the top axis, and ng/mL (the effective concentration measured in the clinical study) along the bottom axis. To convert between µM and ng/mL, we have used that the molecular weight of dabrafenib is 519.6 g/mol^53^.

In order to investigate three-drug vertical inhibition of the BRAF-MEK-ERK pathway in BRAF-V600E mutant melanoma, the model could directly be extended to include the effects of an ERK inhibitor. As a further development of the modelling, one could study the effects of dabrafenib-trametinib treatment scheduling in a multi-cellular system using a multi-scale, agent-based model (ABM)^46,47^. In order to implement the pathway model into an ABM, the relationship between ERK-activity and cell cycle progression/inhibition and cell death needs to be established^32,33,48^. This is one aim of our future modelling studies. ABMs are naturally able to incorporate spatio-temporal variations in intracellular BRAF and ATP concentrations amongst cells, where these variations are derived from both genetic, phenotypic and environmental factors^49,50^. ABMs can thus be used to simulate how drug resistant tumour subclones (*e*.*g*., melanoma cells with elevated BRAF-levels) and drug sensitive tumour subclones evolve in time and space in response to various drug combinations, drug doses and drug treatment schedules. As an alternative future model extension to spatially explicit and stochastic ABMs, the model developed in study can be integrated within a deterministic, age-structured cell population model in which the cell cycle dynamics is dependent on ERK activity.

Thus, via a data-driven and bottom-up modelling approach^51^, the *in silico* model developed in this study can be built upon to simulate how BRAF-MEK-ERK inhibition affects, not only intracellular ERK-activity, but also, the progression of drug resistance in melanoma tumours. Previous experimental work has shown that drug combinations, drug doses, and treatment schedules all impact the evolution of drug resistant tumour subclones in melanoma, when the tumours are subjected to inhibitors that target the BRAF-MEK-ERK pathway^4,52^.

The predictive power of quantitative *in silico* models, such as that described here, are dependent on the accuracy of the kinetic constants for the different components of the signalling cascade as well as on the relative concentrations of the individual enzymes in the cascade. We would suggest that in the light of their key importance further studies are needed to validate the accuracy of the kinetic constants used.

Indeed, translating such models to the treatment of individual cancer patients in the future may require the concentration and activity of the signalling components to be measured on an individual basis.

The mathematical model described in this study provides a method to quantitatively assess how vertical dabrafenib-trametinib inhibition suppresses ERK-activity. We note that the modelling approach outlined in this paper can also be applied to other signalling cascades targeted in cancer treatment. Mathematical oncology models that are developed alongside experiments can be used to systematically motivate which drug combinations, drug doses and treatment schedules warrant experimental investigation^47^.

## Additional Information

### Authors’ contributions

All authors designed the study and contributed to the mathematical model. SH carried out the computational simulations. All authors analysed the data and results. SH drafted the manuscript. All authors revised the manuscript. All authors have approved of the manuscript, and have agreed to be accountable for all aspects of the work presented in the manuscript.

### Data availability

Simulation code is available on the code hosting platform GitHub. Instructions on how to access, run and modify the code are available in the Supplementary Material (SM2).

