## Supplementary Material (SM1-5). for "Quantifying ERK-activity in response to inhibition of the BRAFV600E-MEK-ERK cascade using mathematical modelling"

**References 13**

### SM1 Formulating the system of differential algebraic equations

#### SM1.1 The system of reactions (R.1-R.36)

The system of chemical reactions (R.1-R.36), as listed below, describes signalling dynamics in the intracellular BRAFV600E-MEK-ERK cascade, when a cell is subjected to the BRAF-inhibitor dabrafenib and the MEK-inhibitor trametinib. The cascade and drug actions are schematically illustrated in Figures 1 and 2 in the main manuscript, respectively, and (R.1-R.36) are listed in Figure 3. The dot-notation ($\cdot$) is here used to denote complexes of two or more molecles. Reactions involving dabrafenib and trametinib are respectively marked by single (*) and double (**) stars, and can be omitted if one wishes to study the system in the absence of these drugs.

**System of reactions** (R.1-R.36):

(R.1) $\text{BRAF} + \text{ATP} \underset{d_1}{\overset{a_1}{\rightleftharpoons}} \text{BRAF}\cdot\text{ATP}$

(R.2) $\text{BRAF} + \text{MEK} \underset{d_2}{\overset{a_2}{\rightleftharpoons}} \text{BRAF}\cdot\text{MEK}$

(R.3) $\text{BRAF} + \text{pMEK} \underset{d_2}{\overset{a_2}{\rightleftharpoons}} \text{BRAF}\cdot\text{pMEK}$

(R.4) $\text{BRAF}\cdot\text{ATP} + \text{MEK} \underset{d_2}{\overset{a_2}{\rightleftharpoons}} \text{BRAF}\cdot\text{ATP}\cdot\text{MEK} \xrightarrow{k_{12}} \text{BRAF} + \text{ADP} + \text{pMEK}$

(R.5) $\text{BRAF}\cdot\text{ATP} + \text{pMEK} \underset{d_2}{\overset{a_2}{\rightleftharpoons}} \text{BRAF}\cdot\text{ATP}\cdot\text{pMEK} \xrightarrow{k_{12}} \text{BRAF} + \text{ADP} + \text{ppMEK}$

(R.6) $\text{BRAF}\cdot\text{MEK} + \text{ATP} \underset{d_1}{\overset{a_1}{\rightleftharpoons}} \text{BRAF}\cdot\text{ATP}\cdot\text{MEK} \xrightarrow{k_{12}} \text{BRAF} + \text{ADP} + \text{pMEK}$

(R.7) $\text{BRAF}\cdot\text{pMEK} + \text{ATP} \underset{d_1}{\overset{a_1}{\rightleftharpoons}} \text{BRAF}\cdot\text{ATP}\cdot\text{pMEK} \xrightarrow{k_{12}} \text{BRAF} + \text{ADP} + \text{ppMEK}$

(R.8) $\text{pMEK} + \text{phosph}_1 \underset{d_3}{\overset{a_3}{\rightleftharpoons}} \text{pMEK}\cdot\text{phosph}_1 \xrightarrow{k_3} \text{MEK} + \text{phosph}_1$

(R.9) $\text{ppMEK} + \text{phosph}_1 \underset{d_3}{\overset{a_3}{\rightleftharpoons}} \text{ppMEK}\cdot\text{phosph}_1 \xrightarrow{k_3} \text{pMEK} + \text{phosph}_1$

(R.10)* $\text{BRAF} + \text{DBF} \underset{d_4}{\overset{a_4}{\rightleftharpoons}} \text{BRAF}\cdot\text{DBF}$

(R.11)* $\text{BRAF}\cdot\text{MEK} + \text{DBF} \underset{d_4}{\overset{a_4}{\rightleftharpoons}} \text{BRAF}\cdot\text{MEK}\cdot\text{DBF}$

(R.12)* $\text{BRAF}\cdot\text{pMEK} + \text{DBF} \underset{d_4}{\overset{a_4}{\rightleftharpoons}} \text{BRAF}\cdot\text{pMEK}\cdot\text{DBF}$

(R.13)* $\text{BRAF}\cdot\text{DBF} + \text{MEK} \underset{d_2}{\overset{a_2}{\rightleftharpoons}} \text{BRAF}\cdot\text{MEK}\cdot\text{DBF}$

(R.14)* $\text{BRAF}\cdot\text{DBF} + \text{pMEK} \underset{d_2}{\overset{a_2}{\rightleftharpoons}} \text{BRAF}\cdot\text{pMEK}\cdot\text{DBF}$

(R.15) $\text{ppMEK} + \text{ATP} \underset{d_5}{\overset{a_5}{\rightleftharpoons}} \text{ppMEK}\cdot\text{ATP}$

(R.16) $\text{ppMEK} + \text{ERK} \underset{d_6}{\overset{a_6}{\rightleftharpoons}} \text{ppMEK}\cdot\text{ERK}$

(R.17) $\text{ppMEK} + \text{pERK} \underset{d_6}{\overset{a_6}{\rightleftharpoons}} \text{ppMEK}\cdot\text{pERK}$

(R.18) ppMEK·ATP + ERK $\underset{d_6}{\overset{a_6}{\rightleftharpoons}}$ ppMEK·ATP·ERK $\xrightarrow{k_{56}}$ ppMEK + ADP + pERK

(R.19) ppMEK·ATP + pERK $\underset{d_6}{\overset{a_6}{\rightleftharpoons}}$ ppMEK·ATP·pERK $\xrightarrow{k_{56}}$ ppMEK + ADP + ppERK

(R.20) ppMEK·ERK + ATP $\underset{d_5}{\overset{a_5}{\rightleftharpoons}}$ ppMEK·ATP·ERK $\xrightarrow{k_{56}}$ ppMEK + ADP + ppERK

(R.21) ppMEK·pERK + ATP $\underset{d_5}{\overset{a_5}{\rightleftharpoons}}$ ppMEK·ATP·pERK $\xrightarrow{k_{56}}$ ppMEK + ADP + ppERK

(R.22) pERK + $phosph_2$ $\underset{d_7}{\overset{a_7}{\rightleftharpoons}}$ pERK·$phosph_2$ $\xrightarrow{k_7}$ ERK + $phosph_2$

(R.23) ppERK + $phosph_2$ $\underset{d_7}{\overset{a_7}{\rightleftharpoons}}$ ppERK·$phosph_2$ $\xrightarrow{k_7}$ pERK + $phosph_2$

(R.24)** ppMEK + TMT $\underset{d_8}{\overset{a_8}{\rightleftharpoons}}$ ppMEK·TMT

(R.25)** ppMEK·ATP + TMT $\underset{d_8}{\overset{a_8}{\rightleftharpoons}}$ ppMEK·ATP·TMT

(R.26)** ppMEK·ERK + TMT $\underset{d_8}{\overset{a_8}{\rightleftharpoons}}$ ppMEK·ERK·TMT

(R.27)** ppMEK·pERK + TMT $\underset{d_8}{\overset{a_8}{\rightleftharpoons}}$ ppMEK·pERK·TMT

(R.28)** ppMEK·ERK·ATP + TMT $\underset{d_8}{\overset{a_8}{\rightleftharpoons}}$ ppMEK·ERK·ATP·TMT

(R.29)** ppMEK·pERK·ATP + TMT $\underset{d_8}{\overset{a_8}{\rightleftharpoons}}$ ppMEK·pERK·ATP·TMT

(R.30)** ppMEK·TMT + ATP $\underset{d_5}{\overset{a_5}{\rightleftharpoons}}$ ppMEK·ATP·TMT

(R.31)** ppMEK·TMT + ERK $\underset{d_6}{\overset{a_6}{\rightleftharpoons}}$ ppMEK·ERK·TMT

(R.32)** ppMEK·TMT + pERK $\underset{d_6}{\overset{a_6}{\rightleftharpoons}}$ ppMEK·pERK·TMT

(R.33)** ppMEK·TMT·ERK + ATP $\underset{d_5}{\overset{a_5}{\rightleftharpoons}}$ ppMEK·ERK·ATP·TMT

(R.34)** ppMEK·TMT·pERK + ATP $\underset{d_5}{\overset{a_5}{\rightleftharpoons}}$ ppMEK·pERK·ATP·TMT

(R.35)** ppMEK·TMT·ATP + ERK $\underset{d_6}{\overset{a_6}{\rightleftharpoons}}$ ppMEK·ERK·ATP·TMT

(R.36)** ppMEK·TMT·ATP + pERK $\underset{d_6}{\overset{a_6}{\rightleftharpoons}}$ ppMEK·pERK·ATP·TMT

#### SM1.2 The system of ordinary differential equations (O.1-O.36)

Using the law of mass action, the system of reactions (R.1-R.36) can be formulated in terms of a system of ordinary differential equations (O.1-O.36). We here let $\mathbf{y_{36x1}}$ denote a column vector with 36 elements, where each element corresponds to a signalling molecule concentration so that,

$$
\begin{aligned}
&y(1) = [BRAF],\\
&y(2) = [ATP],\\
&y(3) = [ATP \cdot BRAF],\\
&y(4) = [MEK],
\end{aligned}
$$

$$
\begin{aligned}
y(5) &= [BRAF \cdot MEK], \\
y(6) &= [pMEK], \\
y(7) &= [BRAF \cdot pMEK], \\
y(8) &= [ATP \cdot BRAF \cdot MEK], \\
y(9) &= [ADP], \\
y(10) &= [ATP \cdot BRAF \cdot pMEK], \\
y(11) &= [ppMEK], \\
y(12) &= [phosph_1], \\
y(13) &= [pMEK \cdot phosph_1], \\
y(14) &= [phosph_1 \cdot ppMEK], \\
y(15) &= [DBF], \\
y(16) &= [BRAF \cdot DBF], \\
y(17) &= [BRAF \cdot DBF \cdot MEK], \\
y(18) &= [BRAF \cdot DBF \cdot pMEK], \\
y(19) &= [ATP \cdot ppMEK], \\
y(20) &= [ERK], \\
y(21) &= [ERK \cdot ppMEK], \\
y(22) &= [pERK], \\
y(23) &= [pERK \cdot ppMEK], \\
y(24) &= [ATP \cdot ERK \cdot ppMEK], \\
y(25) &= [ATP \cdot pERK \cdot ppMEK], \\
y(26) &= [ppERK], \\
y(27) &= [phosph_2], \\
y(28) &= [pERK \cdot phosph_2], \\
y(29) &= [phosph_2 \cdot ppERK], \\
y(30) &= [TMT], \\
y(31) &= [TMT \cdot ppMEK], \\
y(32) &= [ATP \cdot TMT \cdot ppMEK], \\
y(33) &= [ERK \cdot TMT \cdot ppMEK], \\
y(34) &= [TMT \cdot pERK \cdot ppMEK], \\
y(35) &= [ATP \cdot ERK \cdot TMT \cdot ppMEK], \\
y(36) &= [ATP \cdot TMT \cdot pERK \cdot ppMEK].
\end{aligned}
$$

Using this notation, the system of ODEs (O.1-O.36) can be formulated as,

$$
\begin{aligned}
\frac{dy(1)}{dt} &= y(1)y(2)(-a_1) + d_1 y(3) + y(1)y(4)(-a_2) + d_2 y(5) + y(1)y(6)(-a_2) + d_2 y(7) \\
&\quad + y(8)k_{12} + y(10)k_{12} + y(8)k_{12} + y(10)k_{12} + y(1)y(15)(-a_4) + d_4 y(16),
\end{aligned} \tag{O.1}
$$

$$
\begin{aligned}
\frac{dy(2)}{dt} &= y(1)y(2)(-a_1) + d_1 y(3) + y(5)y(2)(-a_1) + d_1 y(8) + y(7)y(2)(-a_1) + d_1 y(10) \\
&\quad + y(11)y(2)(-a_5) + d_5 y(19) + y(21)y(2)(-a_5) + d_5 y(24) + y(23)y(2)(-a_5) \\
&\quad + d_5 y(25) + y(31)y(2)(-a_5) + d_5 y(32) + y(33)y(2)(-a_5) + d_5 y(35) \\
&\quad + y(34)y(2)(-a_5) + d_5 y(36),
\end{aligned} \tag{O.2}
$$

$$
\frac{dy(3)}{dt} = y(3)(-d_1) + a_1(y(1)y(2)) + y(3)y(4)(-a_2) + d_2 y(8) + y(3)y(6)(-a_2) + d_2 y(10), \tag{O.3}
$$

$$
\begin{aligned}
\frac{dy(4)}{dt} &= y(1)y(4)(-a_2) + d_2 y(5) + y(3)y(4)(-a_2) + d_2 y(8) + y(13)k_3 + y(16)y(4)(-a_2) \\
&\quad + d_2 y(17),
\end{aligned} \tag{O.4}
$$

$$
\frac{dy(5)}{dt} = y(5)(-d_2) + a_2(y(1)y(4)) + y(5)y(2)(-a_1) + d_1 y(8) + y(5)y(15)(-a_4) + d_4 y(17), \tag{O.5}
$$

$$\frac{dy(6)}{dt} = y(1)y(6)(-a_2) + d_2y(7) + y(8)k_{12} + y(3)y(6)(-a_2) + d_2y(10) + y(8)k_{12} + y(6)y(12)(-a_3) + d_3y(13) + y(14)k_3 + y(16)y(6)(-a_2) + d_2y(18), \tag{O.6}$$

$$\frac{dy(7)}{dt} = y(7)(-d_2) + a_2(y(1)y(6)) + y(7)y(2)(-a_1) + d_1y(10) + y(7)y(15)(-a_4) + d_4y(18), \tag{O.7}$$

$$\frac{dy(8)}{dt} = y(8)(-d_2 - k_{12}) + a_2(y(3)y(4)) + y(8)(-d_1 - k_{12}) + a_1(y(5)y(2)), \tag{O.8}$$

$$\frac{dy(9)}{dt} = y(8)k_{12} + y(10)k_{12} + y(8)k_{12} + y(10)k_{12} + y(24)k_{56} + y(25)k_{56} + y(24)k_{56} + y(25)k_{56}, \tag{O.9}$$

$$\frac{dy(10)}{dt} = y(10)(-d_2 - k_{12}) + a_2(y(3)y(6)) + y(10)(-d_1 - k_{12}) + a_1(y(7)y(2)), \tag{O.10}$$

$$\frac{dy(11)}{dt} = y(10)k_{12} + y(10)k_{12} + y(11)y(12)(-a_3) + d_3y(14) + y(11)y(2)(-a_5) + d_5y(19) + y(11)y(20)(-a_6) + d_6y(21) + y(11)y(22)(-a_6) + d_6y(23) + y(24)k_{56} + y(25)k_{56} + y(24)k_{56} + y(25)k_{56} + y(11)y(30)(-a_8) + d_8y(31), \tag{O.11}$$

$$\frac{dy(12)}{dt} = y(6)y(12)(-a_3) + d_3y(13) + y(13)k_3 + y(11)y(12)(-a_3) + d_3y(14) + y(14)k_3, \tag{O.12}$$

$$\frac{dy(13)}{dt} = y(13)(-d_3 - k_3) + a_3(y(6)y(12)), \tag{O.13}$$

$$\frac{dy(14)}{dt} = y(14)(-d_3 - k_3) + a_3(y(11)y(12)), \tag{O.14}$$

$$\frac{dy(15)}{dt} = y(1)y(15)(-a_4) + d_4y(16) + y(5)y(15)(-a_4) + d_4y(17) + y(7)y(15)(-a_4) + d_4y(18), \tag{O.15}$$

$$\frac{dy(16)}{dt} = y(16)(-d_4) + a_4(y(1)y(15)) + y(16)y(4)(-a_2) + d_2y(17) + y(16)y(6)(-a_2) + d_2y(18), \tag{O.16}$$

$$\frac{dy(17)}{dt} = y(17)(-d_4) + a_4(y(5)y(15)) + y(17)(-d_2) + a_2(y(16)y(4)), \tag{O.17}$$

$$\frac{dy(18)}{dt} = y(18)(-d_4) + a_4(y(7)y(15)) + y(18)(-d_2) + a_2(y(16)y(6)), \tag{O.18}$$

$$\frac{dy(19)}{dt} = y(19)(-d_5) + a_5(y(11)y(2)) + y(19)y(20)(-a_6) + d_6y(24) + y(19)y(22)(-a_6) + d_6y(25) + y(19)y(30)(-a_8) + d_8y(32), \tag{O.19}$$

$$\frac{dy(20)}{dt} = y(11)y(20)(-a_6) + d_6y(21) + y(19)y(20)(-a_6) + d_6y(24) + y(28)k_7 + y(31)y(20)(-a_6) + d_6y(33) + y(32)y(20)(-a_6) + d_6y(35), \tag{O.20}$$

$$\frac{dy(21)}{dt} = y(21)(-d_6) + a_6(y(11)y(20)) + y(21)y(2)(-a_5) + d_5y(24) + y(21)y(30)(-a_8) + d_8y(33), \tag{O.21}$$

$$\frac{dy(22)}{dt} = y(11)y(22)(-a_6) + d_6y(23) + y(24)k_{56} + y(19)y(22)(-a_6) + d_6y(25) + y(24)k_{56} + y(22)y(27)(-a_7) + d_7y(28) + y(29)k_7 + y(31)y(22)(-a_6) + d_6y(34) + y(32)y(22)(-a_6) + d_6y(36), \tag{O.22}$$

$$\frac{dy(23)}{dt} = y(23)(-d_6) + a_6(y(11)y(22)) + y(23)y(2)(-a_5) + d_5y(25) + y(23)y(30)(-a_8) + d_8y(34), \tag{O.23}$$

$$\frac{dy(24)}{dt} = y(24)(-d_6 - k_{56}) + a_6(y(19)y(20)) + y(24)(-d_5 - k_{56}) + a_5(y(21)y(2)) + y(24)y(30)(-a_8) + d_8y(35), \quad (O.24)$$

$$\frac{dy(25)}{dt} = y(25)(-d_6 - k_{56}) + a_6(y(19)y(22)) + y(25)(-d_5 - k_{56}) + a_5(y(23)y(2)) + y(25)y(30)(-a_8) + d_8y(36), \quad (O.25)$$

$$\frac{dy(26)}{dt} = y(25)k_{56} + y(25)k_{56} + y(26)y(27)(-a_7) + d_7y(29), \quad (O.26)$$

$$\frac{dy(27)}{dt} = y(22)y(27)(-a_7) + d_7y(28) + y(28)k_7 + y(26)y(27)(-a_7) + d_7y(29) + y(29)k_7, \quad (O.27)$$

$$\frac{dy(28)}{dt} = y(28)(-d_7 - k_7) + a_7(y(22)y(27)), \quad (O.28)$$

$$\frac{dy(29)}{dt} = y(29)(-d_7 - k_7) + a_7(y(26)y(27)), \quad (O.29)$$

$$\frac{dy(30)}{dt} = y(11)y(30)(-a_8) + d_8y(31) + y(19)y(30)(-a_8) + d_8y(32) + y(21)y(30)(-a_8) + d_8y(33) + y(23)y(30)(-a_8) + d_8y(34) + y(24)y(30)(-a_8) + d_8y(35) + y(25)y(30)(-a_8) + d_8y(36), \quad (O.30)$$

$$\frac{dy(31)}{dt} = y(31)(-d_8) + a_8(y(11)y(30)) + y(31)y(2)(-a_5) + d_5y(32) + y(31)y(20)(-a_6) + d_6y(33) + y(31)y(22)(-a_6) + d_6y(34), \quad (O.31)$$

$$\frac{dy(32)}{dt} = y(32)(-d_8) + a_8(y(19)y(30)) + y(32)(-d_5) + a_5(y(31)y(2)) + y(32)y(20)(-a_6) + d_6y(35) + y(32)y(22)(-a_6) + d_6y(36), \quad (O.32)$$

$$\frac{dy(33)}{dt} = y(33)(-d_8) + a_8(y(21)y(30)) + y(33)(-d_6) + a_6(y(31)y(20)) + y(33)y(2)(-a_5) + d_5y(35), \quad (O.33)$$

$$\frac{dy(34)}{dt} = y(34)(-d_8) + a_8(y(23)y(30)) + y(34)(-d_6) + a_6(y(31)y(22)) + y(34)y(2)(-a_5) + d_5y(36), \quad (O.34)$$

$$\frac{dy(35)}{dt} = y(35)(-d_8) + a_8(y(24)y(30)) + y(35)(-d_5) + a_5(y(33)y(2)) + y(35)(-d_6) + a_6(y(32)y(20)), \quad (O.35)$$

$$\frac{dy(36)}{dt} = y(36)(-d_8) + a_8(y(25)y(30)) + y(36)(-d_5) + a_5(y(34)y(2)) + y(36)(-d_6) + a_6(y(32)y(22)). \quad (O.36)$$

#### SM1.3 The conservation laws (C.1-C.7)

The regarded system described by (O.1-O.36) obeys a set of conservation laws (C.1-C.7) which are,

$$0 = y(1) + y(3) + y(5) + y(7) + y(8) + y(10) + y(16) + y(17) + y(18) - BRAF_{tot}, \quad (C.1)$$

$$0 = y(4) + y(5) + y(6) + y(7) + y(8) + y(10) + y(11) + y(13) + y(14) + y(17) + y(18) + y(19) + y(21) + y(23) + y(24) + y(25) + y(31) + y(32) + y(33) + y(34) + y(35) + y(36) - MEK_{tot}, \quad (C.2)$$

$$0 = y(20) + y(21) + y(22) + y(23) + y(24) + y(25) + y(26) + y(28) + y(29) + y(33) + y(34) + y(35) + y(36) - ERK_{tot}, \quad (C.3)$$

$$0 = y(2) + y(3) + y(8) + y(10) + y(19) + y(24) + y(25) + y(32) + y(35) + y(36) - ATP_{tot}, \quad \text{(C.4)}$$
$$0 = y(15) + y(16) + y(17) + y(18) - DBF_{tot}, \quad \text{(C.5)}$$
$$0 = y(30) + y(31) + y(32) + y(33) + y(34) + y(35) + y(36) - TMT_{tot}, \quad \text{(C.6)}$$
$$0 = y(12) + y(13) + y(14) - phosph_{1,tot}, \quad \text{(C.7)}$$
$$0 = y(27) + y(28) + y(29) - phosph_{2,tot}. \quad \text{(C.8)}$$

Note that, we assume a cellular ADP-to-ATP recycling to continually occur and thus the $[ADP]$ variable is not included in the $[ATP]$ conservation law. This ATP-to-ADP recycling is not explicitly included in the model. In C.1-C.7, the total concentrations, denoted by a $tot$-subscript, are equal to the initial concentrations described in Table ST2 (Section SM3.2) so that,

$$\begin{aligned} BRAF_{tot} &= \text{[BRAF](0)}, \\ MEK_{tot} &= \text{[MEK](0)}, \\ ERK_{tot} &= \text{[ERK](0)}, \\ phosph_{1,tot} &= [\text{phosph}_1](0), \\ phosph_{2,tot} &= [\text{phosph}_2](0), \\ DBF_{tot} &= \text{[DBF](0)}, \\ TMT_{tot} &= \text{[TMT](0)}. \end{aligned}$$

### SM2 Code access and instructions

The computational MATLAB [1] code includes functions to convert the system of reactions (R.1-R.36) into a system of differential algebraic equations, *i.e.,* (O.1-O.36) with code-generated substitutions (C.1-C.7), using the law of mass action. The MATLAB function `ode15s` is then used to solve for $\mathbf{y_{36x1}}$ (*i.e.*, the signalling molecule concentration vector) as a function of system concentrations and time.

#### SM2.1 Code access - GitHub repository

The MATLAB code, developed for this project, is available on the code hosting platform GitHub, in the project repository (https://github.com/SJHamis/MAPKcascades).

#### SM2.2 Instructions on how to run the code

The model outputs/result plots presented in the main manuscript can be generated by running the five main-files, with file names `main_[insert option].m`. The code includes three sub-directories: (1) `auxiliary_files_plots`, which includes files needed to generate data for specific time points, (2) `auxiliary_files_model_setup`, which includes files needed to set up the DAE MATLAB function file (`mapk_cascade_DAE.m`), (3) `model_details`, in which the system of reactions (R.1-R.36) and the rate constants are manually set.

#### SM2.3 Instructions on how to modify the code

Instructions on how to edit the cascade structure, rate constants and initial conditions are provided below. For more detailed information, see in-line comments in the MATLAB code.

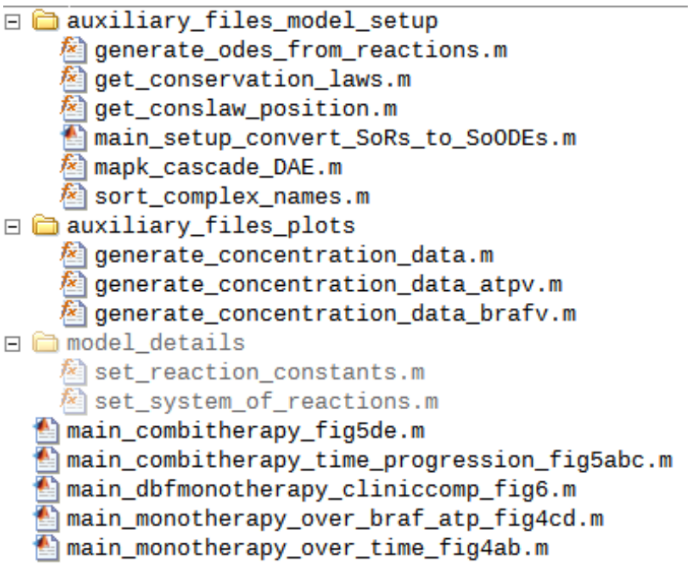

Figure SF1: Code directories.

##### SM2.3.1 Modifying the rate constants

The rate constants can be manually edited in the file
`model_details/set_reaction_constants.m`.
By first editing the constants and thereafter running the file
`auxiliary_files_model_setup/main_setup_convert_SoRs_to_SoODEs.m`,
the MATLAB DAE function file
`auxiliary_files_model_setup/mapk_cascade_DAE.m`
will be updated to include the modified rate constants.

##### SM2.3.2 Modifying the chemical reactions/cascade structure

The system of reactions can be manually edited in the file
`model_details/set_system_of_reactions.m`.
By first editing the system of reactions, by for example by including more terms, and thereafter running the file
`auxiliary_files_model_setup/main_setup_convert_SoRs_to_SoODEs.m`,
the function files
`auxilary_files_model_setup/mapk_cascade_DAE.m`
and
`auxilary_files_model_setup/get_conslaw_position.m`
will be updated to include the new model equations (ODEs and conservation laws).

##### SM2.3.3 Modifying the initial conditions

The model initial condition (*i.e.,* the initial cascade component concentrations) can be modified in the main files,
`main_[insert option].m`.

### SM3 Model parameters

#### SM3.1 Kinetic constants

| Rate constants in the system of reactions (R.1-R.36) | | | | |
|---|---|---|---|---|
| constant | value | reference | description | appearance |
| $a_1$ | $0.106\mu M^{-1}s^{-1}$ | [2, 3] | Forward rate constant for ATP binding to BRAF (or a BRAF complex with MEK/pMEK). | R.1,R.6,R.7 |
| $d_1$ | $6.23\ s^{-1}$ | [2, 3, 4] | Reverse rate constant for ATP binding to BRAF (or a BRAF complex with MEK/pMEK). | R.1,R.6,R.7 |
| $a_2$ | $0.106\mu M^{-1}s^{-1}$ | † | Forward rate constant for MEK/pMEK binding to BRAF (or a BRAF complex with ATP/DBF). | R.2,R.3,R.4, R.5,R.13,R.14 |
| $d_2$ | $0.02385\ s^{-1}$ | [2, 3] | Reverse rate constant for MEK/pMEK binding to BRAF (or a BRAF complex with ATP/DBF). | R.2,R.3,R.4, R.5,R.13,R.14 |
| $k_{1,2}$ | $0.66\ s^{-1}$ | [3] | Catalytic rate constant for production of pMEK/ppMEK from BRAF·ATP complex with MEK/pMEK. | R.4,R.5,R.6, R.7 |
| $a_3$ | $0.106\mu M^{-1}s^{-1}$ | † | Forward rate constant for phosph$_1$ binding to pMEK/ppMEK. | R.8,R.9 |
| $d_3$ | $0.0159\ s^{-1}$ | [2, 3] | Reverse rate constant for phosph$_1$ binding to pMEK/ppMEK. | R.8,R.9 |
| $k_3$ | $0.0159\ s^{-1}$ | [2, 3] | Catalytic rate constant for production of MEK/pMEK from phosph$_1$ complex with pMEK/ppMEK. | R.8,R.9 |
| $a_4$ | $0.106\mu M^{-1}s^{-1}$ | † | Forward rate constant for the drug DBF binding to BRAF (or a BRAF complex with MEK/pMEK). | R.10,R.11,R.12 |
| $d_4$ | $0.0000593\ s^{-1}$ | [2, 3, 5, 6, 7, 8] | Reverse rate constant for the drug DBF binding to BRAF (or a BRAF complex with MEK/pMEK). | R.10,R.11,R.12 |
| $a_5$ | $0.106\mu M^{-1}s^{-1}$ | † | Forward rate constant for ATP binding to ppMEK (or a ppMEK complex with ERK/pERK/TMT/TMT&ERK/TMT&pERK). | R.15,R.20,R.21, R.30,R.33,R.34 |
| $d_5$ | $0.3468\ s^{-1}$ | [2, 3, 9] | Reverse rate constant for ATP binding to ppMEK (or a ppMEK complex with ERK/pERK/TMT/TMT&ERK/TMT&pERK). | R.15,R.20,R.21, R.30,R.33,R.34 |
| $a_6$ | $0.106\mu M^{-1}s^{-1}$ | † | Forward rate constant for ERK/pERK binding to ppMEK (or a ppMEK complex with ATP/TMT/TMT&ATP). | R.16,R.17,R.18, R.19,R.31,R.32, R.35,R.36 |
| $d_6$ | $0.01184\ s^{-1}$ | [2, 3, 9] | Reverse rate constant for ERK/pERK binding to ppMEK (or a ppMEK complex with ATP/TMT/TMT&ATP). | R.16,R.17,R.18, R.19,R.31,R.32, R.35,R.36 |
| $k_{5,6}$ | $0.0242\ s^{-1}$ | [9] | Catalytic rate constant for production of pERK/ppERK from ppMEK·ATP complex with ERK/pERK. | R.18,R.19,R.20, R.21 |
| $a_7$ | $0.106\mu M^{-1}s^{-1}$ | † | Forward rate constant for phosph$_2$ binding to pERK/ppERK. | R.22,R.23 |
| $d_7$ | $0.0159\ s^{-1}$ | [2, 3] | Reverse rate constant for phosph$_2$ binding to pERK/ppERK. | R.22,R.23 |
| $k_7$ | $0.0159\ s^{-1}$ | [2, 3] | Catalytic rate constant for production of ERK/pERK from phosph$_2$ complex with pERK/ppERK. | R.22,R.23 |
| $a_8$ | $0.106\mu M^{-1}s^{-1}$ | † | Forward rate constant for the drug TMT binding to ppMEK (or a ppMEK complex with ATP/ERK/pERK/ERK&ATP/pERK&ATP). | R.24,R.25,R.26, R.27,R.28,R.29 |
| $d_8$ | $0.0012296\ s^{-1}$ | [2, 3, 10] | Reverse rate constant for the drug TMT binding to ppMEK (or a ppMEK complex with ATP/ERK/pERK/ERK&ATP/pERK&ATP). | R.24,R.25,R.26, R.27,R.28,R.29 |

Table ST1: A table of the rate constants appearing in the system of reactions (R.1-R.36). The system includes eight forward rate constants $a_1, a_2, .., a_8$, eight reverse rate constants $d_1, d_2, .., d_8$, and four catalytic rate constants $k_{1,2}, k_3, k_{5,6}, k_7$. Stroke symbols (/) in the description column denote 'or', whilst ampersand symbols (&) denote 'and'. The rightmost column describes in which of the reactions (R.1-R.36) the constants appear. † All forward rate constants are of the same value.

#### SM3.2 Model initial condition (initial cascade component concentrations)

| Initial concentrations | | |
|---|---|---|
| Initial condition | value | reference |
| [BRAF](0) | 3-10 nM* | [2] |
| [MEK](0) | 1.2 $\mu$M | [2] |
| [ERK](0) | 1.2 $\mu$M | [2] |
| [phosph1](0) | 0.3 nM | [2] |
| [phosph2](0) | 0.12 $\mu$M | [2] |
| [ATP](0) | 1-5 mM** | [9] |
| [DBF](0) | *applied DBF dose* (here 0-10 $\mu$M) | |
| [TMT](0) | *applied TMT dose* (here 0-10 $\mu$M) | |

Table ST2: Initial model concentrations, where the (0) notation denotes ‘at time zero’. For molecule concentrations not listed in this table, the initial conditions is 0 $\mu$M. *Huang & Ferrell [2] listed MAPKKK=3nM, and we use this as our baseline value. However, in order to study amplified BRAF levels as a mode of drug resistance, we explore initial BRAF concentrations within the range 3-10 nM. Similarly, initial ATP concentrations are explored within the range 1-5mM, but the baseline value is 1mM.

### SM4 Data and derivation of model parameters

All model parameter values for the forward rate constants $a_1, a_2, ..., a_8$, the reverse rate constants $d_1, d_2, ..., d_8$ and the catalytic rate constant $k_{1,2}, k_3, k_{5,6}, k_7$ are obtained from data that is available in the literature. Note that $a_i = a_1 \forall i = 2, 3, ...8$, as is explained in the main manuscript (in **Methods - Model parameters**).

#### SM4.1 Rate constants for BRAF-ATP and BRAF-MEK reactions

VanScyoc *et al.* [3] provide kinetic data for the phosphorylation of the substrate MEK by wild type BRAF. We use their data to approximate the parameter values for $a_1$, $d_2$ and $k_{1,2}$ in our model, *i.e.* rate constants for the BRAF-ATP and BRAF-MEK reactions. We also compute the reverse BRAF-ATP rate constant for wild type BRAF, here denoted by $d_{1,WT}$. We then use $d_{1,WT}$ to find $a_1$, as the catalytic rate constant $k_{1,2}$ and the forward rate constant $a_1$, can be assumed to be the same for both V600E-mutant and wild type BRAF interacting with ATP. In order to set $d_1$, we thereafter use data for $\text{BRAF}^{\text{V600E}}$-ATP interactions provided by Hatzivassiliou *et al.* [4].

**BRAF-ATP reactions** (approximating $a_1, d_1, k_{1,2}$): VanScyoc *et al.* report that the disassociation constant for ATP binding to wild type BRAF is $K_{d1,WT}(\text{ATP}) = 7.8 \pm 2.2\mu\text{M}$. We use this $K_d$-value to find the relationship between $a_1$ and $d_{1,WT}$ using,

$$K_{d1,WT} = \frac{d_{1,WT}}{a_1} \approx 7.8\mu\text{M} \Longrightarrow d_{1,WT} \approx 7.8\mu\text{M} \cdot a_1. \tag{P.1}$$

VanScyoc *et al.* further report the catalytic rate constant for BRAF-ATP reactions to be $k_{cat} = 0.66 \pm 0.30\text{s}^{-1}$, which we can directly use as a parameter value for $k_{1,2}$ so that

$$k_{1,2} \approx 0.66\text{s}^{-1}. \tag{P.2}$$

We approximate the Michaelis constant, $K_{m1,WT}$, as the average value of $K_m$(ATP) at 0.15 and 0.6 $\mu$M MEK1, reported by VanScyoc *et al.*, so that $K_{m1,WT} \approx 14\mu\text{M}$. We can now use (P.1), (P.2) and the data for the Michaelis constant to solve for the parameter value of $a_1$,

$$K_{m1,WT} = \frac{d_{1,WT} + k_{1,2}}{a_1} \Longrightarrow a_1 \cdot K_{m1,WT} = d_{1,WT} + k_{1,2} \Longrightarrow a_1 \cdot 14\mu\text{M} \approx (7.8\mu\text{M} \cdot a_1) + 0.66\text{s}^{-1} \Longrightarrow$$

$$\Longrightarrow a_1(14\mu\text{M} - 7.8\mu\text{M}) \approx 0.66\text{s}^{-1} \Longrightarrow a_1 \cdot 6.2\mu\text{M} \approx 0.66\text{s}^{-1} \Longrightarrow a_1 \approx \frac{0.66\text{s}^{-1}}{6.2\mu\text{M}} = 0.106\text{s}^{-1}(\mu\text{M})^{-1}. \quad \text{(P.3)}$$

If we now plug this value of $a_1$ into the expression for $d_{1,WT}$ in (P.1) we obtain,

$$d_{1,WT} \approx 7.8\mu\text{M} \cdot a_1 = 7.8\mu\text{M} \cdot 0.106\text{s}^{-1}(\mu\text{M})^{-1} = 0.8268\text{s}^{-1}. \quad \text{(P.4)}$$

In order to now find $d_1$, the reverse rate constant for V600E mutant BRAF, we use the ATP-BRAF$^{\text{V600E}}$ $K_m$ value reported by Hatzivassiliou *et al.* which is 65 $\mu$M. Using this, we can obtain $d_1$ via

$$K_{m,1} = \frac{d_1 + k_{1,2}}{a_1} \Longrightarrow d_1 = K_{m,1}a_1 - k_{1,2} = 65\mu\text{M} \cdot 0.106\text{s}^{-1}(\mu\text{M})^{-1} - 0.66\text{s}^{-1} = 6.23\text{s}^{-1}. \quad \text{(P.5)}$$

Hatzivassiliou *et al.* further report that the ATP-BRAF $K_m$ value is 5$\mu$M for wild type BRAF, which is in good agreement with VanScyoc *et al.*'s data.

**BRAF-MEK reactions** (approximating $d_2$): From VanScyoc *et al.*'s data, we can approximate the disassociation constant $K_{d2} = d_2/a_2$ as $K_{0.5}$(MEK1) $\approx 0.225\mu$M, where this value is the average of the reported $K_{0.5}$ values at 10 and 100 $\mu$M ATP. As $a_2 = a_1$, we can now find $d_2$ via

$$K_{0.5} = K_{d2} = \frac{d_2}{a_2} \approx 0.225\mu\text{M} \Longrightarrow d_2 \approx 0.225\mu\text{M} \cdot a_2 = 0.225\mu\text{M} \cdot a_1 \approx \quad \text{(P.6)}$$
$$\approx 0.225\mu\text{M} \cdot 0.106s^{-1}(\mu\text{M})^{-1} = 0.02385s^{-1}.$$

Note that the catalytic rate constant for BRAF-MEK, and ATP, reactions was obtained in (P.2).

#### SM4.2 Rate constants for MEK-ATP and MEK-ERK reactions

Mansour *et al.* [9] provide steady-state rate constants for MEK reactions. We use data from their study to set model parameter values describing MEK-ATP and MEK-ERK reactions.

**MEK-ATP interactions** (approximating $d_5, k_{5,6}$): Mansour *et al.* report that $V_m = 1.45 \pm 0.05\text{min}^{-1}$. After re-expressing this value in units of 'per second', we can directly use this data to approximate our parameter value of $k_{5,6}$,

$$k_{5,6} \approx 0.0242\text{s}^{-1}. \quad \text{(P.7)}$$

Mansour *et al.* further report that $K_{m,ATP} = 3.5 \pm 0.8\mu$M, which corresponds to $K_{m5}$ in our model so that,

$$K_{m5} = \frac{d_5 + k_{5,6}}{a_5} \approx 3.5\mu\text{M} \Longrightarrow \frac{d_5 + 0.0242\text{s}^{-1}}{a_5} \approx 3.5\mu\text{M} \Longrightarrow d_5 \approx 3.5\mu\text{M} \cdot a_5 - 0.0242\text{s}^{-1}, \quad \text{(P.8)}$$

where we have used (P.7). By further using the convention that $a_5 = a_1 \approx 0.106s^{-1}(\mu\text{M})^{-1}$, we obtain a value for $d_5$ such that,

$$d_5 \approx 3.5\mu\text{M} \cdot a_1 - 0.0242\text{s}^{-1} \approx 3.5\mu\text{M} \cdot 0.106\text{s}^{-1}(\mu\text{M})^{-1} - 0.0242\text{s}^{-1} = 0.3468\text{s}^{-1}. \quad \text{(P.9)}$$

**MEK-ERK interactions** (approximating $d_6$): From Mansour *et al.*, the Michaelis constant for $K_{m6}$ can be found, as $K_{m,ERK2} = 0.34 \pm 0.06\mu$M and thus

$$K_{m6} = \frac{d_6 + k_{5,6}}{a_6} \approx 0.34\mu\text{M}. \quad \text{(P.10)}$$

As $a_6 = a_1$ and $k_{5,6}$ have already been set, we can now solve for $d_6$,

$$\frac{d_6 + k_{5,6}}{a_1} \approx 0.34\mu\text{M} \Longrightarrow \frac{d_6 + 0.0242\text{s}^{-1}}{0.106s^{-1}(\mu\text{M})^{-1}} \approx 0.34\mu\text{M} \Longrightarrow d_6 \approx 0.34\mu\text{M} \cdot 0.106s^{-1}(\mu\text{M})^{-1} - 0.0242\text{s}^{-1} = \tag{P.11}$$

$$= 0.01184\text{s}^{-1}.$$

#### SM4.3 Rate constants for phosphatase reactions

**phosph$_1$ and phosph$_1$ reactions** (approximating $d_3, k_3, d_7, k_7$): Huang & Ferrell [2] estimated that, for all reactions,

$$K_m = \frac{d+k}{a} = 0.3\mu\text{M}. \tag{P.12}$$

We use this expression for the phosphatase reactions $i = 3$ and $i = 7$, so that

$$d_{3/7} + k_{3/7} = 0.3\mu\text{M} \cdot a_{3/7}, \tag{P.13}$$

where the slash notation (/) denotes *'or'*. By further estimating that $d_3 = k_3$ and $d_7 = k_7$, we obtain

$$d_{3/7} = k_{3/7} = \frac{0.3\mu\text{M} \cdot a_{3/7}}{2} = 0.15\mu\text{M} \cdot a_{3/7}, \tag{P.14}$$

and with $a_{3/7} = a_1$ we get,

$$d_{3/7} = k_{3/7} = 0.15\mu\text{M} \cdot a_1 \approx 0.15\mu\text{M} \cdot 0.106\text{s}^{-1}(\mu\text{M})^{-1} \approx 0.0159\text{s}^{-1}. \tag{P.15}$$

#### SM4.4 Rate constants for drug reactions

**BRAF-DBF interactions** (approximating $d_4$): Data pertaining to DBF reactions is gathered from the US Food and Drug Administration [5], according to which the *in vitro* IC50 value is $0.65$nM. Rheault *et al.* [6] similarly report this IC50 value to be $0.7$nM. We use this IC50 value to estimate the inhibitory constant $K_i$ via the Cheng-Prusoff approach so that,

$$K_i = \frac{\text{IC50}}{1 + [L]/K_d} = \frac{0.65\text{nM}}{1 + 10\mu\text{M}/65\mu\text{M}} = 0.56\text{nM}, \tag{P.16}$$

where the experimental ligand (ATP) concentration is $10\mu$M [7], and the ATP $K_d$ value has been approximated as $K_m = 65\mu$M, as previously used. We can now find $d_4$ via,

$$K_{i4} = \frac{d_4}{a_4} = 0.00056\mu\text{M} \Longrightarrow d_4 = 0.00056\mu\text{M} \cdot a_4. \tag{P.17}$$

With $a_4 = a_1 \approx 0.106\text{s}^{-1}(\mu\text{M})^{-1}$, the value of $d_4$ can now be obtained as

$$d_4 = 0.00056\mu\text{M} \cdot a_1 \approx 0.00056\mu\text{M} \cdot 0.106\text{s}^{-1}(\mu\text{M})^{-1} = 5.93 \cdot 10^{-5}s^{-1}. \tag{P.18}$$

Koelblinger *et al.* [8] report the reverse rate constant for dabrafenib-BRAF interactions to be $9.6 \cdot 10^{-5}s^{-1}$, which is in good agreement with our calculated value.

**MEK-TMT interactions** (approximating $d_8$): TMT data is gathered from Gilmartin *et al.* [10] who report that $K_i = 0.0116 \pm 0.0007\mu$M. With the median as our $K_{d8}$ value, we obtain

$$K_{d8} = \frac{d_8}{a_8} \Longrightarrow d_8 = 0.0116\mu\text{M} \cdot a_8 = 0.0116\mu\text{M} \cdot a_1 \approx 0.0116\mu\text{M} \cdot 0.106\text{s}^{-1}(\mu\text{M})^{-1} = 0.0012296\text{s}^{-1}. \tag{P.19}$$

### SM5 Results not included in main manuscript

#### SM5.1 Extended time range dabrafenib responses

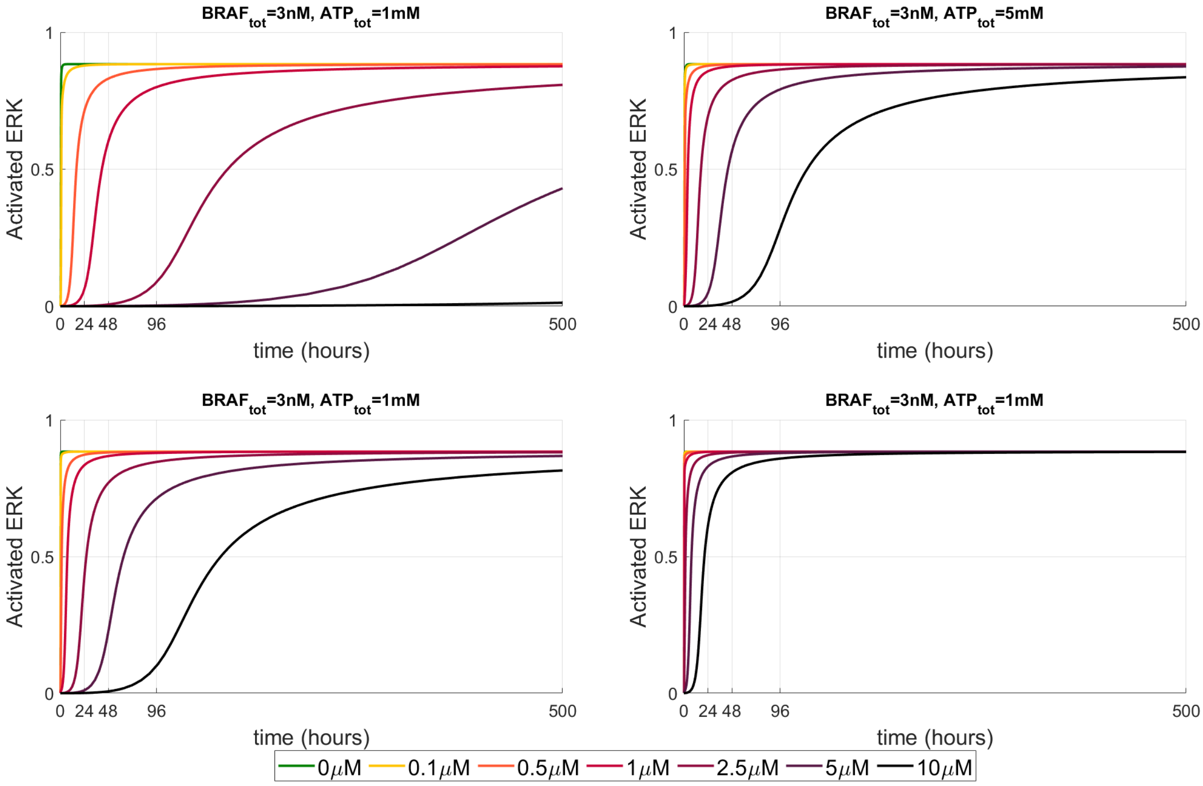

Figure SF2: Modified version of Figure 4a in the main manuscript, with a time span of 0 to 500 hours. Dabrafenib monotherapy doses are indicated by the legend at the bottom of the Figure.
